# PDAC-ANN: an artificial neural network to predict Pancreatic Ductal Adenocarcinoma based on gene expression

**DOI:** 10.1101/698209

**Authors:** Palloma Porto Almeida, Cristina Padre Cardoso, Leandro Martins de Freitas

## Abstract

**Background:** Although the pancreatic ductal adenocarcinoma (PDAC) presents high mortality and metastatic potential, there is a lack of effective therapies and a low survival rate for this disease. This PDAC scenario urges new strategies for diagnosis, drug targets, and treatment.

**Methods:** We performed a gene expression microarray meta-analysis of the tumor against healthy tissues in order to identify differentially expressed genes shared among all datasets, named core-genes (CG). We confirmed the pancreatic expressed proteins of the CG through The Human Protein Atlas. The five most expressed proteins in the tumor group were selected to train an artificial neural network to classify samples.

**Results:** This microarray included 110 tumor and 77 healthy samples. We identified a CG composed of 60 genes, 58 upregulated and two downregulated. The upregulated CG included proteins and extracellular matrix receptors linked to actin cytoskeleton reorganization. With the Human Protein Atlas, we verified that thirteen genes of the CG are translated, with high or medium expression in most of the pancreatic tumor samples. To train our artificial neural network, we used the five most expressed genes (KRT19, LAMC2, MELK, MET, TOP2A). The artificial neural network model (PDAC-ANN) classified the train samples with sensitivity of 0.95, specificity of 0.9, and f1-score of 0.93. The PDAC-ANN could classify the test samples with a sensitivity of 0.97, specificity of 0.88, and f1-score 0.94.

**Conclusion:** The gene expression meta-analysis and confirmation of the protein expression allow us to select five genes highly expressed PDAC samples. We could build a python script to classify the samples based on mRNA expression. This software can be useful in the PDAC diagnosis.

## Background

The pancreatic ductal adenocarcinoma (PDAC) is the most common pancreatic cancer histological subtype with high mortality due to the lack of symptoms in the initial phase of the disease and its aggressive progression [1,2]. PDAC development is directly related to *KRAS* overexpression [2, 3], along with the inactivation of the tumor-suppressor genes *CDKN2A/p16* [4], *SMAD4/DPC4* [5] and *TP53* [6, 7]. The *KRAS* activation is considered significant in PDAC progression, and many efforts were made to inhibit its activity [8]; nevertheless, it seems to be undruggable [9]. Data have been presented in the literature over integrated analysis about PDAC genes and proteins, classifying PDAC in different molecular subtypes among patients [10], and through integrated genome analyses that reinforce the participation of *KRAS, TP53, SMAD4*, and *CDKN2A* in a subset of PDAC tumors [11].

Since there is a lack of effective therapies and a low survival rate, the research for new biomarkers and therapies targets in PDAC remains active [12–14]. There are some gene expression changes in pancreatic cancer already described and presented as biological markers. The genes in the ribosome and the spliceosome pathway (ribosomal protein genes Nup170, Nup160, and HNRNPU) were described as potential biomarkers [15]. The meta-analysis of PDAC microarray data could identify five biomarkers (TMPRSS4, AHNAK2, POSTN, ECT2, SERPINB5) that classified the PDAC and normal samples with sensitivity of 94%, and specificity of 89.6% [16].

Advances in high-performance computing, such as system biology and artificial intelligence (AI) allows integration of data and pattern recognition that generates not only new understating about diseases but also support new targets discovery and biomarkers development for future treatments [17]. The potential to classify the cancer samples using gene expression, methylation information, and AI has been used in other types of cancers studies and with promising results. The application of these studies would improve the classification of the samples in tumors diagnosis and subtyping [18–20]. The studies using automatic technics to predict risk/diagnosis had demonstrated a high classification performance, presenting sensitivity >90%[21–24].

The high number of features coming from microarray gene expression and methylation genomic information used to train AI tumor diagnosis models can give good results in the classification of samples [18, 19] lowering the false negative rate in training and validation samples. However, the high number of features can make the diagnosis available only for samples with thousands of gene expression values [18]. It has been shown that reducing the number of features can give the same or better results than using thousands of features[25, 26]

The application of AI in pancreatic tumor must improve the early diagnostic and consequently, the treatment and patient survival. The AI has been used to predict risk/diagnosis using pancreatic image and personal health features [27]. The prediction of pancreatic cancer risk in patients with type 2 diabetes was compared using logistic regression and artificial neural network (ANN), again using personal health features and presenting the performance of models predicting the cancer risk factor [24]□. There are also AI models to diagnosis pancreatic cancer-based in four plasma protein selected in mass spectra, showing the potential of AI in predicting the status of the sample based in biological markers with high sensitivity (90.9%) and specificity (91.1%)[22]. The Lustgarten Foundation, created to pancreatic cancer research, pointed the importance of including the AI in the PDAC diagnosis based on magnetic resonance imaging and computed tomography scans [28]. The use of new technologies to help pancreatic cancer risk/diagnosis must be pursued and it would improve patients’ survival. The gene expression changes in pancreatic cancer could be used as biological markers and help in the diagnosis and be used to build a computational model using AI to predict samples status.

In this paper, we performed a meta-analysis of gene expression of public microarray data. We identified a core-genes (CG) group and accessed the protein expression through the Protein Atlas database based in immunohistochemical staining images. Clusterization methods were applied to distinguish between normal and PDAC samples. It was selected five genes combining microarray expression and Protein Atlas information. The gene expression information from PDAC and healthy samples were used to build an ANN (PDAC-ANN). The PDAC-ANN uses gene expression information to predict the sample status (normal or PDAC) and give the probability of the sample be PDAC. This is the first-time gene expression is used to build an ANN model to predict PDAC diagnosis. The results showed here must be verified in a large sample and could be used in the discrimination of samples using these markers. This PDAC-ANN is free software and could be used to improve the diagnosis and help PDAC patients.

## Methods

### Dataset acquisition

The microarray expression data of human healthy and pancreatic cancer tissue were collected from Gene Expression Omnibus (GEO) (https://www.ncbi.nlm.nih.gov/geo/) using the search term “pancreatic ductal adenocarcinoma” and selecting mRNA expression profiling by array. The seven datasets (Table 1) were selected following the criteria: inclusion of (1) studies presenting tumor/non-tumor samples from PDAC; exclusion of studies (2) with induced mutations or activated pathways; (3) cells previously exposed to chemotherapy drugs. These criteria ensure that the expression alterations were provided only from the shift healthy to disease, and not due to induced mutations in cell lineage or chemotherapy treatment. The datasets were loaded into the R software [29] using the GEOquery package [30]. A six studies were analyzed to find differentially expressed genes (DEG), and the seventh microarray study served as an independent dataset to validate the CG derived from the meta-analysis.

**Table 1.** Characteristics of studies used in the meta-analysis.

| Accession number | Study | Array platform | Differentially expressed genes |  | Samples |  |
| --- | --- | --- | --- | --- | --- | --- |
|  |  |  | Upregulated | Downregulated | Tumor | Control |
| GSE23397 | * | Affymetrix Human Exon 1.0 ST Array | 4031 | 870 | 15 | 6 |
| GSE28735 | [65]; [66] | Affymetrix Human Gene 1.0 ST Array | 245 | 146 | 45 | 45 |
| GSE32676 | [67] | Affymetrix Human Genome U133 Plus 2.0 Array | 686 | 319 | 25 | 7 |
| GSE41368 | [68] | Affymetrix Human Gene 1.0 ST Array | 1200 | 462 | 6 | 6 |
| GSE43795 | [69] | Illumina HumanHT-12 V4.0 expression beadchip | 1978 | 1343 | 6 | 5 |
| GSE71989 | * | Affymetrix Human Genome U133 Plus 2.0 Array | 3052 | 661 | 13 | 8 |
| Total |  |  |  |  | **110** | **77** |
| GSE16515 (Validation data set) | [70] | Affymetrix Human Genome U133 Plus 2.0 Array | - | - | 36 | 16 |

\* No publication available
- Analysis of differentially expressed genes was not applied to the validation dataset

### Data processing

Non-specific filtering was applied to each dataset coming from the same GEO series. Briefly, the package genefilter were used to remove the genes with no expression variation among samples [31], followed by the collapse of multiple probe measurements of a given gene into a single gene measurement in package WGCNA [32]. The limma package [33] was used to identify the DEG through a t-test. We considered DEGs when log2 fold change (logFC) was ≥ to 1 and adjusted p-value by false discovery rate ≤ 0.05 [34, 35].

### Core-gene analysis

The DEG frequency among the microarray studies was retrieved, and those shared by all microarray studies were considered as the CG. The CG gene expression values were standardized applying the method 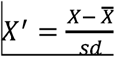, where 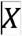 represents the expression values, 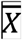 the gene expression average, and 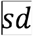 standard deviation [36]. This standardization was followed by a min-max data rescale, for each gene transforming all values to [0, 1] range. Thus, restricting values from different studies to the same range [37]. The CG standardized values were used in the Principal Component Analysis (PCA) and the hierarchical clustering in order to check the clustering of the samples from all datasets based on the CG expression values.

### Data Validation

The immunohistochemical staining images and the protein expression data from pancreatic cancer tissue were used as validation of the CG. Protein expression data were obtained from The Human Protein Atlas (THPA) (www.proteinatlas.org) [38]. We search CG protein expression in pancreatic cancer, and then the expression information was classified as high, medium, low, and not detected. We also investigated a validation using the CG standardized values in a seventh independent dataset (GSE16515). We applied the hierarchical clustering/heatmap and the PCA to the GSE16515 samples to evaluate its capability to differentiate tumor and non-tumor groups.

### Neural Network Sample Classification

We build an ANN using python to classify the sample in healthy or tumor samples. The ANN was trained using normalized gene expression values [0, 1] from the five most expressed genes among the six datasets. We explore the performances of 90 networks architecture with one input layer with five nodes, one or two hidden layers varying the number of nodes from 2 to 10, and two output nodes. Each network architecture was trained 30 times, and we took the mean accuracy in the train set to evaluate the classification performance. We used a learning rate of 0.2, 200 epochs during training, tanh and softmax as activation functions for internal and output node, respectively. The network weights were randomly initialized with values between [-1, 1], and bias with value 1.

## Results

### Differentially expressed genes in meta-analysis

To profile differentially expressed genes in PDAC, we performed a meta-analysis of microarray data available in GEO (Table 1). We collected and compared 110 tumor samples to 77 healthy tissues. We have identified 7821 unique DEG, where 5717 were upregulated and 2104 downregulated genes (logFC = 1; adj. p-value ≤ 0.05) (Additional file 1: Table S1), exhibiting a CG of 60 genes, where 58 were upregulated, and two downregulated (Table 2).

**Table 2.**
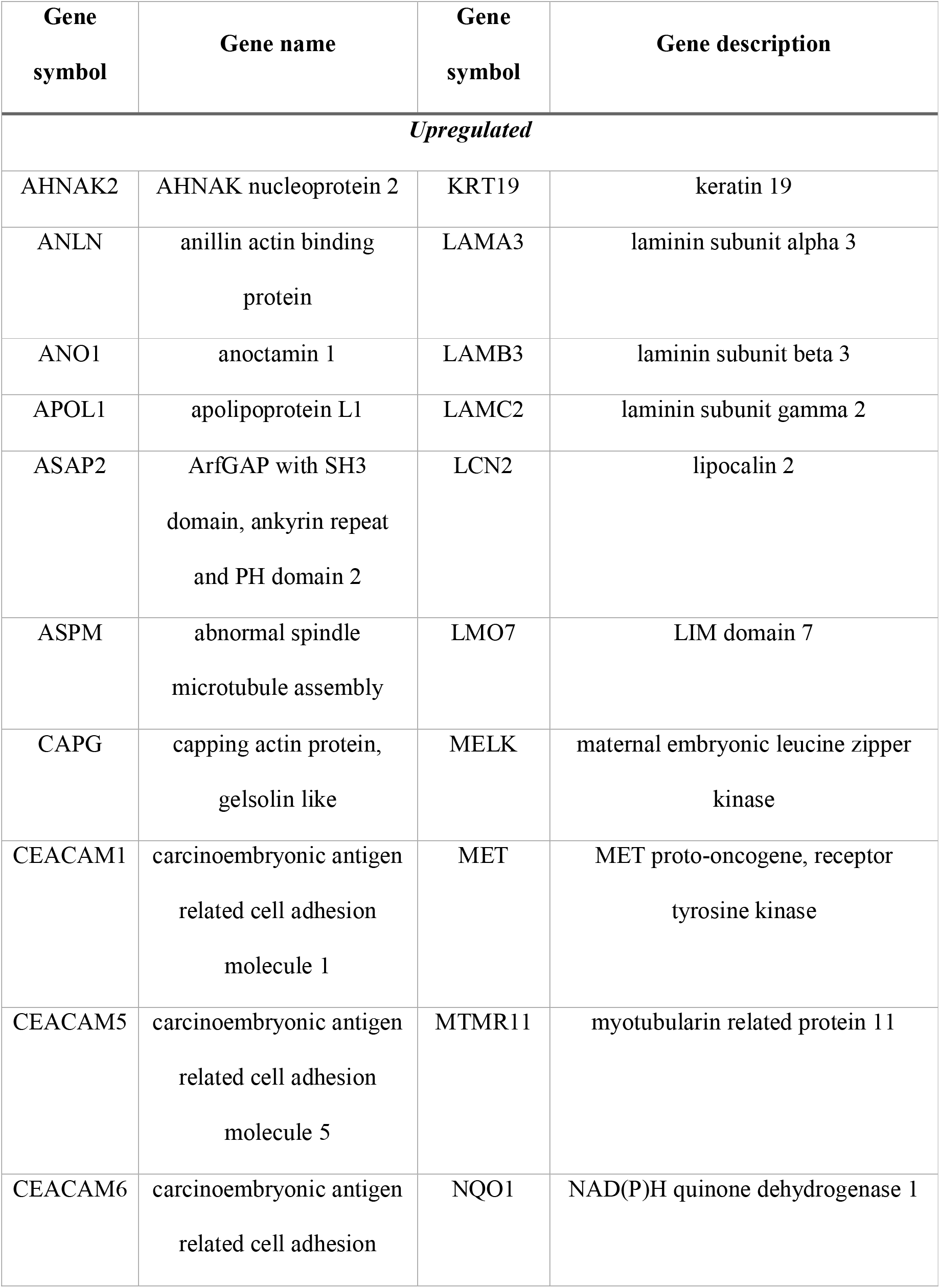
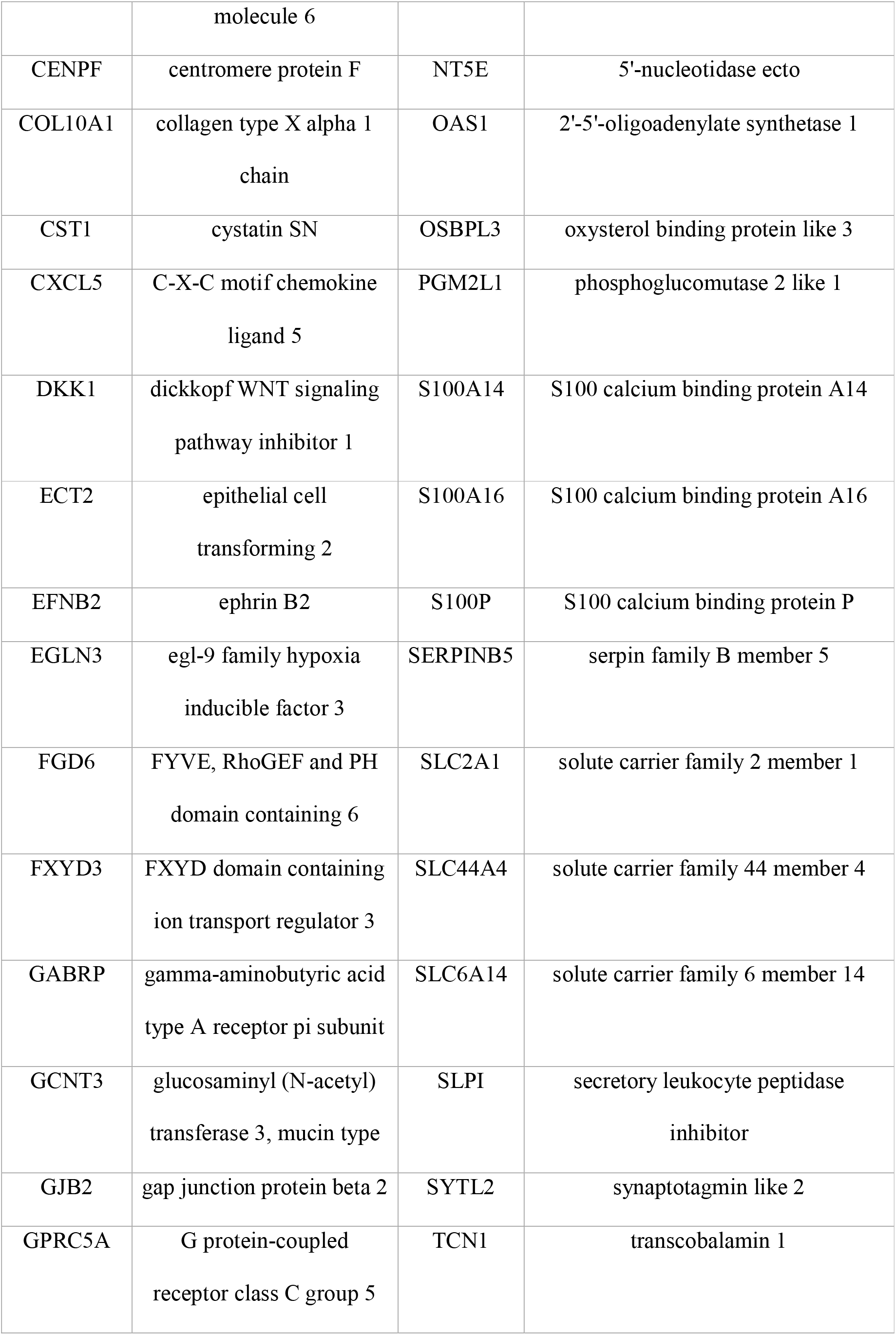
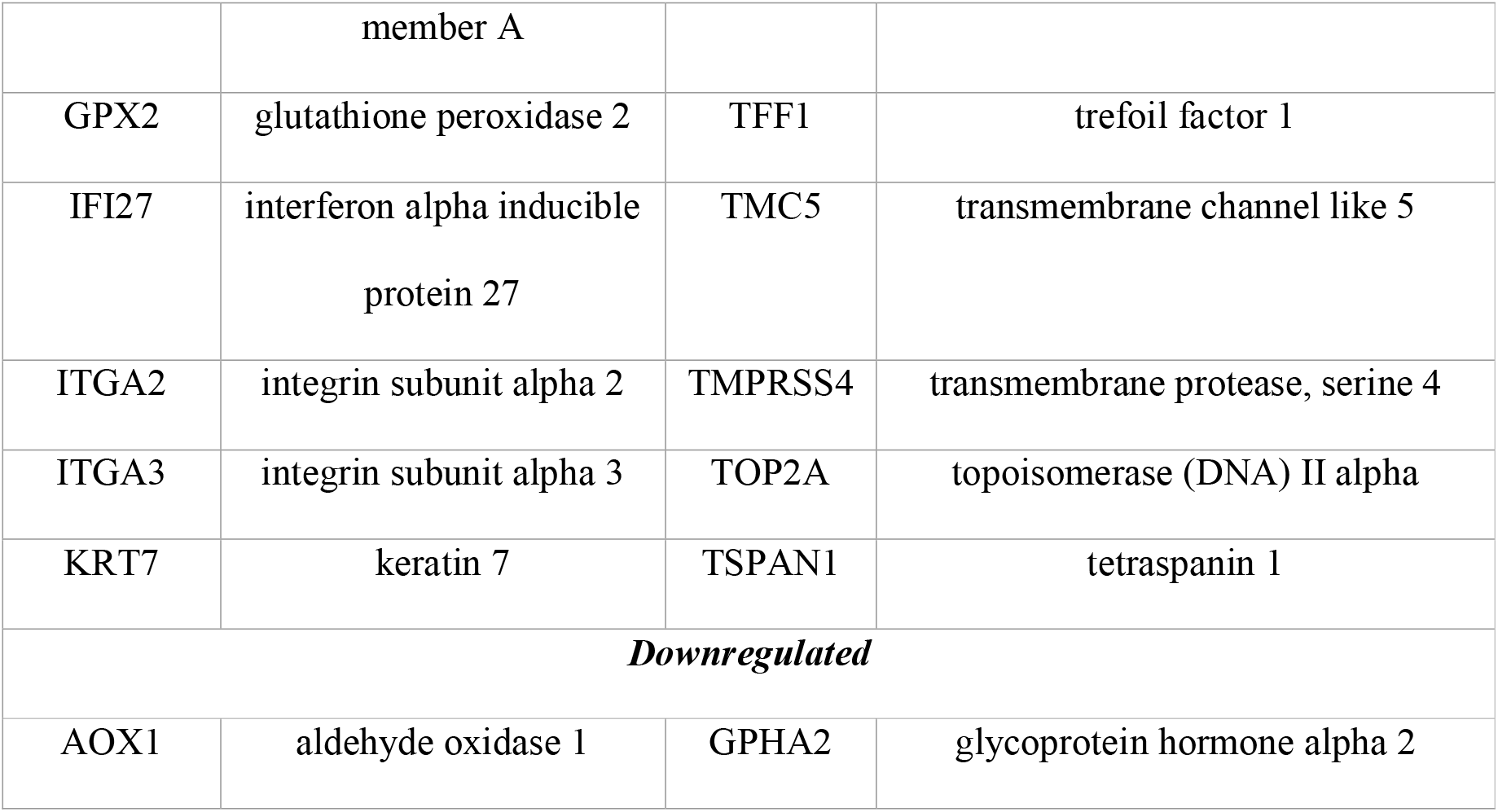
Description of the core-genes involved in PDAC biological process.

The CG showed a profile of upregulated genes functions related to cell membrane-ECM interaction (*LAMA3*, *LAMB3, LAMC2*), cytoskeleton interaction/calcium management *(GCNT3, ANLN, S100A14-16, S100P*), and structural integrity of epithelial cells (*ITGA2, ITGA3, KRT7, KRT19*). Most of the genes reinter the importance of the ECM interaction and cellular morphology in carcinogenic processes in PDAC. The AOX1 and GPHA2 were the only two downregulated genes in PDAC samples compared to control samples. The AOX1 was already detected as downregulated in other PDAC studies [39, 40], and this corroborates the result presented here.

### Immunohistochemical staining images validation

To determine whether the CG is also present as proteins expressed in PDAC, we investigated the expression of these genes in THPA. This analysis could confirm the protein expression from many of the CG list using information from immunohistochemical staining images. The protein expression data from the CG list showed that 31 genes have more than 50% of images with high or medium expression in pancreatic cancer (Fig 2).

**Figure 1.**
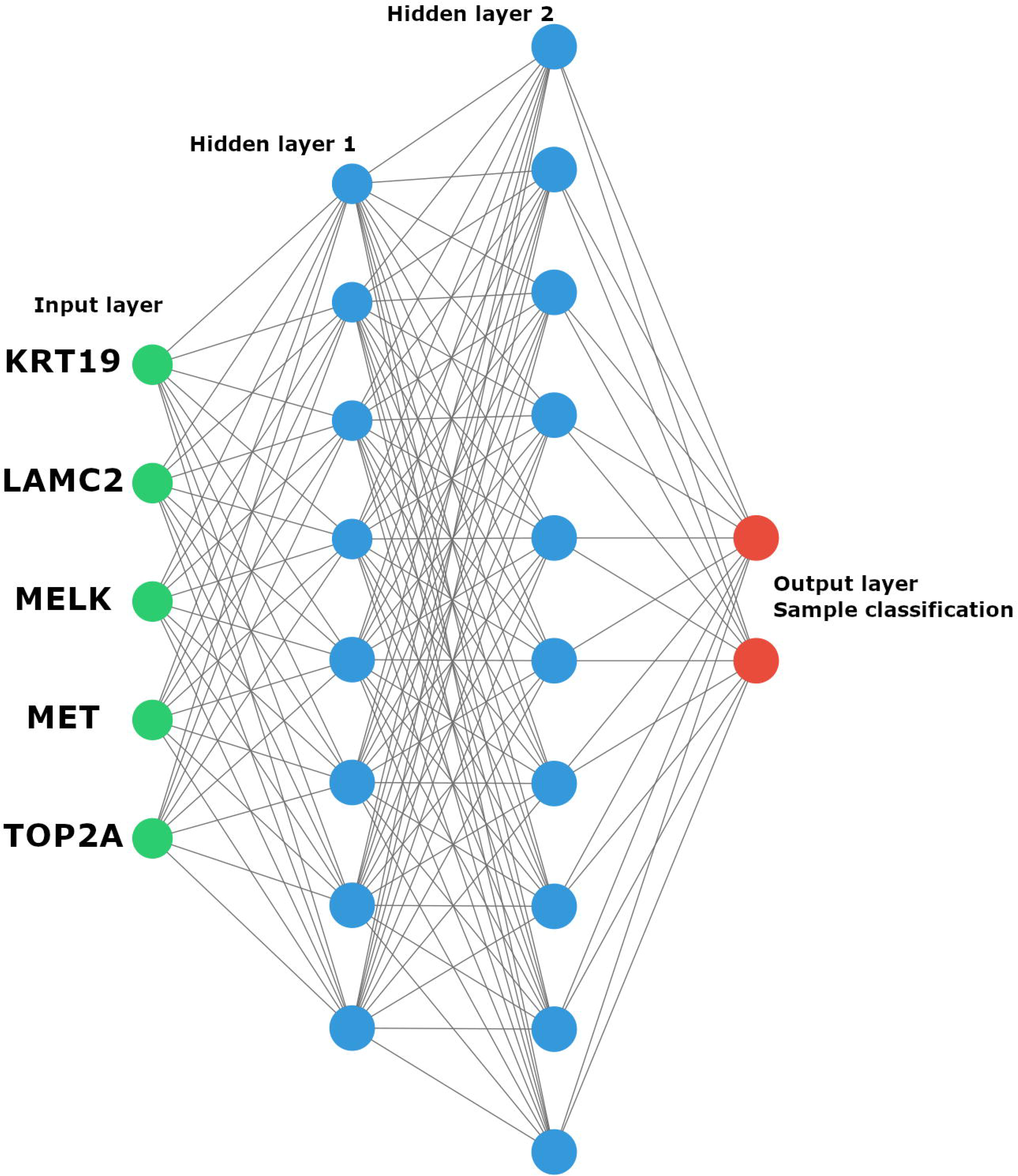
The Artificial neural network architecture. The KRT19, LAMC2, MELK, MET, and TOP2A expression values are data from the input layer (green neurons), The hidden layers (blue neurons) process the expression values, and the output layer (red neurons) give the likelihood of being normal and PDAC sample.

**Fig 2.**
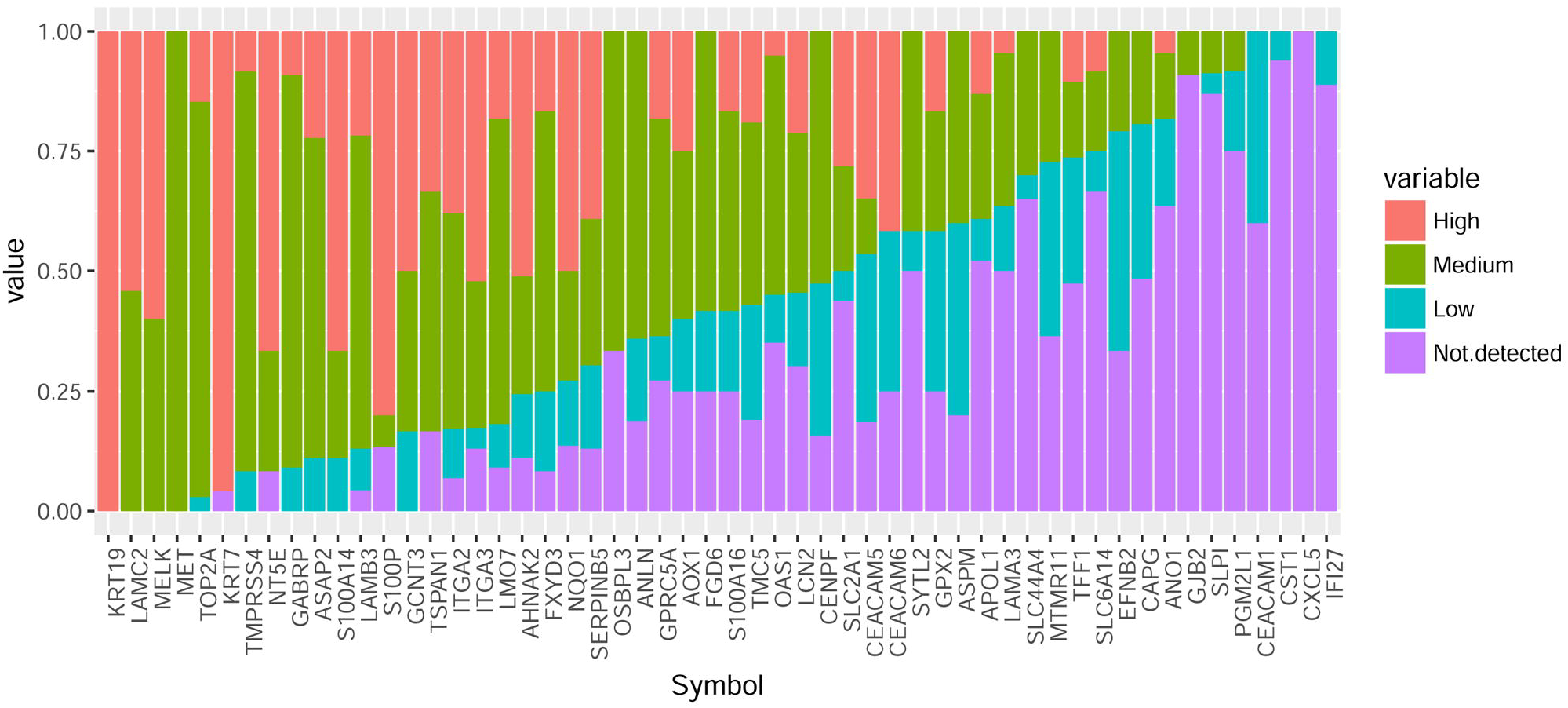
Variation in protein expression data from the GC list retrieved from immunohistochemical staining images in THPA. The protein expression data shows that 31 genes have more than 50% of images with high or medium expression in pancreatic cancer, evidencing the expression of predicted core-genes in the pancreatic tissue — data credit: THPA.

There were three proteins (KRT19, KRT7, and S100P) with more than 75% of immunohistochemical staining images representing high expression values, from a set of 23, 23, 12 images, respectively (Fig 3). The proteins LAMC2, MELK, TOP2A, and MET are also among the set of proteins with the highest expression from the CG list since these proteins showed only high or medium expression levels. The genes IFI27, CXCL5, CST1, and CEACAM1, have a low or not detected expression, not corroborating with the CG list. The protein AOX1 presents a different expression between the mRNA and protein level. The AOX1 protein is highly expressed in some samples (60%) and low or not detected in others (40%).

**Fig 3.**
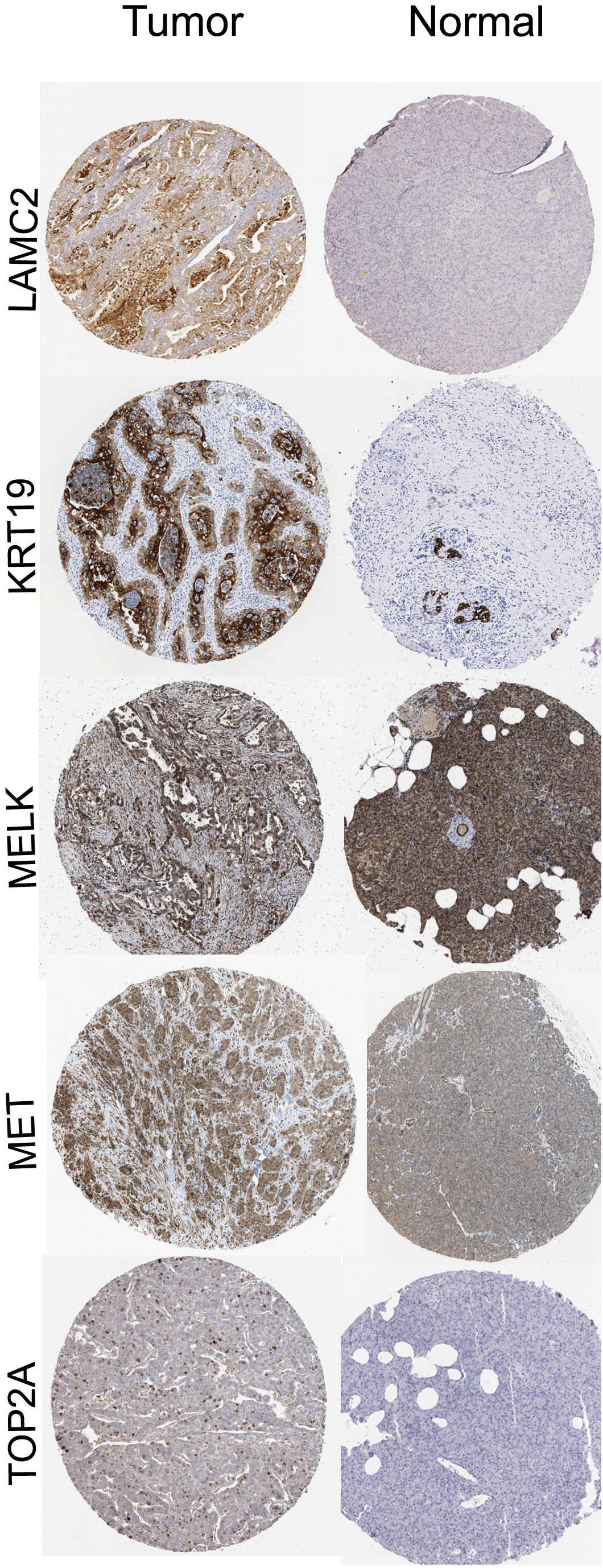
Representative immunohistochemistry staining of LAMC2, KRT19, MELK, MET, and TOP2A in Pancreatic Ductal Adenocarcinoma (Tumor) and normal pancreatic tissue (Normal). The proteins presented more than 75% of images with high expression. Scales bars represent 200 μm — image credit: THPA.

There were six proteins (COL10A1, DKK1, ECT2, EGLN3, TCN1, GPHA2) with no information in THPA, thus is not possible to say about the protein expression in pancreatic cancer. All these data show essential genes in PDAC highly expressed in proteins level, confirming 31 genes from the CG in pancreatic cancer.

### Classification of the merged samples in tumor and control using PCA and hierarchical clustering

We performed hierarchical clustering of the samples/genes and a PCA analysis of the samples to evaluate how different the gene expression is among the samples and how the samples cluster. The PCA showed variation in the expression in a continuous manner, and some PDAC samples mixed with healthy samples. The PCA plot has a region with only PDAC samples almost, indicating more specific gene expression in PDAC. The PCA result indicates a difference in the CG expression enough to classify the samples in healthy and PDAC, although the PCA does not predict the label of the sample (Additional file 2: Figure S1).

The hierarchical clustering, performed using CG expression standardized values from all six datasets, reveal the presence of two groups, and it is possible to check the error of the sample classification (Fig 4). The standardized CG expression values were able to classify the data into two groups, once more indicating that these groups exhibit distinctly cellular processes and functions. The hierarchical clustering showed the ratio Normal Classified/Normal = 0.92 and Tumor Classified/Tumor = 0.83.

**Fig 4.**
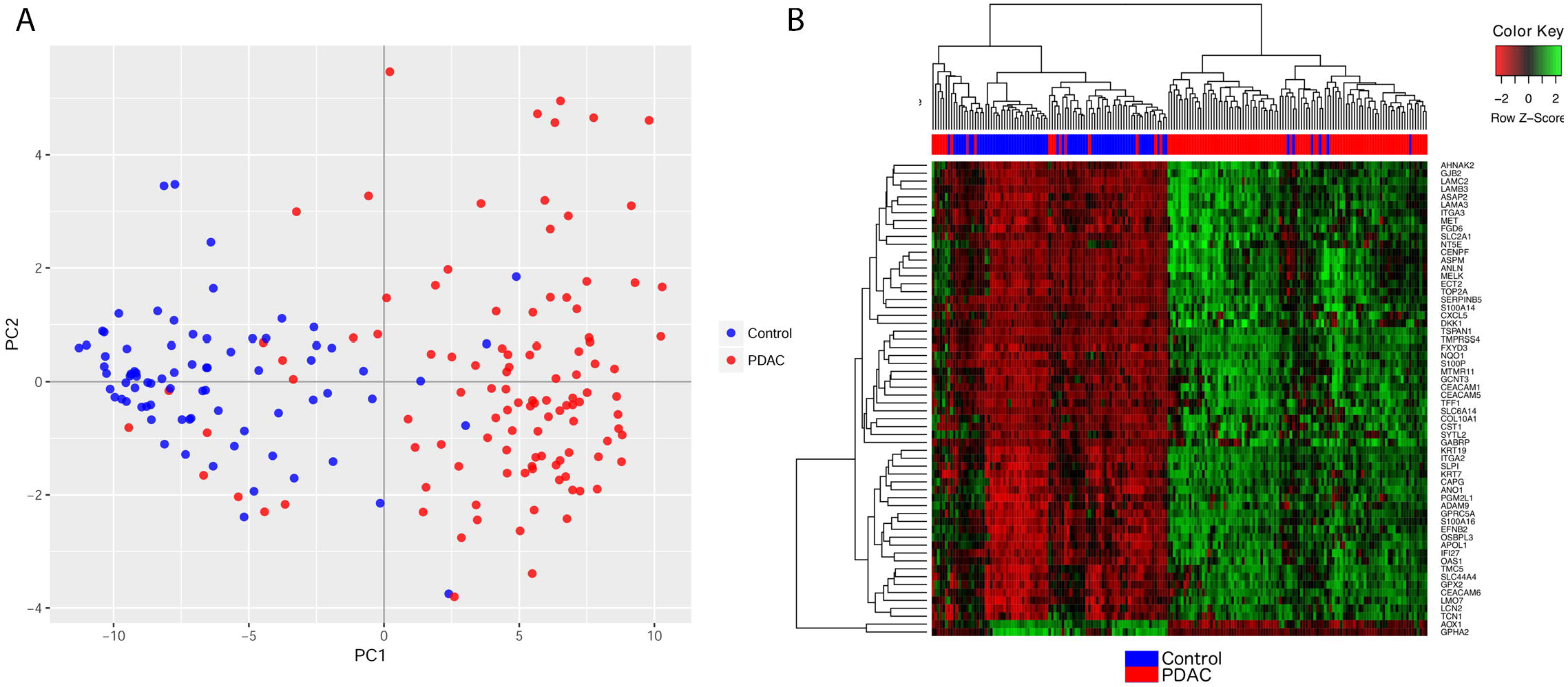
PCA and hierarchical analysis of the merged data set into one data. A. PCA analysis clearly showed two distinct groups corresponding to normal and tumor samples. B. Clustering analysis. Red band indicates the PDAC samples with similar gene expression on 60-core-gene and blue band indicates the normal samples.

The methodology was also applied to an independent dataset (GSE16515) to validate the CG founded by the meta-analysis. The CG expression values from GSE16515 could produce similar results in both PCA and heatmap hierarchical clustering analysis (Figure S1.), showing that CG can classify the data in two groups of normal and tumoral samples, suggesting the CG maps central process in PDAC. Together, these results indicate that the CG expression can distinguish the groups normal from PDAC samples, showing different functional/cellular processes expressed by this condition, and this points to CG list as critical genes in PDAC that could be used to classify the samples and improve diagnosis.

### Neural network sample classification

The best neural network architecture had an accuracy of 93.04 and 94.23% in the train and test set respectively; the architecture has five input neurons, eight and ten neurons in the next two hidden layers, and one output. We examined the classification performance in the validation dataset using the f1-score (0.94), that summarize the precision and recall measurements (table 3). The PDAC-ANN model predicted the trains samples status and achieved sensitivity and specificity of 0.95 and 0.9 respectively; while the prediction of the test samples achieved sensitivity and specificity of 0.88 and 0.97, respectively. The confusion matrix showed that the number of true negative (normal) was 14/16, while the number of true positive is 35/36 (table 4).

**Table 3.** Classification report of the validation test set.

|  | **Precision** | **Recall** | **F1-score** | **Support** |
| --- | --- | --- | --- | --- |
| **Normal** | 0.93 | 0.88 | 0.9 | 16 |
| **Tumor** | 0.95 | 0.97 | 0.96 | 36 |
| **Avg/total** | 0.94 | 0.94 | 0.94 | 52 |

**Table 4.** Confusion matrix of the training and validation test samples.

|  |  | **Actual normal** | **Actual tumor** |
| --- | --- | --- | --- |
| **Training** | **Classified normal** | 69 | 5 |
|  | **Classified tumor** | 8 | 105 |
|  |  | Specificity = 0.9 | Sensitivity = 0.95 |
| **Test** | **Classified normal** | 14 | 1 |
|  | **Classified tumor** | 2 | 35 |
|  |  | Specificity = 0.88 | Sensitivity = 0.97 |

## Discussion

We performed a meta-analysis of mRNA expression data recovered from public datasets deposited in GEO, intending to investigate the profile of molecular alterations in pancreatic ductal adenocarcinoma and use this information to build an ANN predictor. Comparing 110 tumor samples to 77 normal tissues, we were able to observe a central group of genes linked to carcinogenic processes, labeled core-genes, further investigated the protein expression with immunohistochemistry information recovery from THPA, and validated with a seventh microarray through hierarchical clustering, and PCA.

The late diagnosis and high mortality rate in PDAC patients demand better tools to auxiliary in the diagnosis. Currently, the gold standard blood-based biomarker for PDAC diagnosis is the CA 19-9 [41]. However, CA 19-9 lacks the sensitivity for the early detection and also has a poor predictive value in asymptomatic patients [42–44]. Thus, the precise selection of biomarkers can increase the accuracy in the diagnosis of PDAC as well as provide a cheaper diagnostic method with a lower invasion.

We performed a validation of the CG through the immunohistochemical images retrieved from THPA. Our results indicate a list of possible biomarkers to PDAC. Furthermore, we present two biomarkers differentially expressed and often used for PDAC diagnosis, carcinoembryonic antigen-related cell adhesion molecule 5 (CEACAM5, also known as CEA) and Apolipoprotein B (ApoB). The ApoB has already been described as a carrier for CA 19-9 antigen [29], and the CEACAM5 has been pointed as the second serum biomarker most used clinically for detecting PDAC [28]. We confirmed the expression of thirty-one genes from CG with high expression in the protein level. These proteins are involved in many functions in cancer biology. For instance, the most expressed protein, keratin 19 (KRT19), is a structural protein of epithelial cells, with expression in a subset of pancreatic cells [45]. The KRT19 was already described as a possible biomarker for PDAC and patients with upregulation of KRT19 presents poor differentiation, large tumor size, lymph node metastasis, and invasion [46]. In other gastrointestinal cancers, clinical pathological analyses revel KRT19 correlated with metastasis, tumor size, microvascular invasion, decreased tumor differentiation, and also conferred an invasive phenotype [46].

The laminin subunit gamma 2 (LAMC2) protein involved in cell survival was shown to be upregulated in PDAC samples using microarray and immunohistochemical analyses, and a biomarker for diagnosis and prognosis integrating a multigene panel [47]. Proteomic analysis pointed the LAMC2 as a potential biomarker for PDAC, being upregulated with an mRNA fold of 8.36. The serum concentration of LAMC2 in patients with PDAC was ∼ 3.5-fold higher from benign and healthy samples, indicating this gene as a promising biomarker [48]. PDAC patients expressing high amount of LAMC2 have a poor prognosis [49], reinforcing this gene as a putative biomarker for diagnosis and prognosis.

The maternal embryonic leucine zipper kinase (MELK) is expressed in stem/progenitor cell populations, including hematopoietic stem cells, but also in a variety of cancers [50, 51]. Its function has been implicated in cell cycle regulation [52] and cell division [53]. MELK expression was associated with therapeutic resistance and poor prognosis [54], and thus, reported as a promising molecular target for cancer therapy [55]. In adults, MELK regulates the size and number of regenerative pancreatic ducts, promoting cell migration in cancer, with no action in adult β cell regeneration [56]. The function of MELK in PDAC is unknown; however, the MELK inhibitor, OTS167, presented preclinical efficacy with antitumor activity in both hematological and solid cancer, including pancreatic cancer xenografic models [57].

The metastatic spreading of PDAC requires heterogeneous stromal signaling related to MET proto-oncogene, receptor tyrosine kinase (MET) and its ligand, hepatocyte growth factor (HGF). The MET/HGF axis have been associated with poor overall survival of pancreatic cancer patients [58]. MET gene is also involved in resistance to palliative chemotherapy in PDAC patients, possibly due to the downstream pathway of MET/HGF axis in the maintenance of pancreatic progenitor cells and stem cells [59]. Therefore, MET/HGF pathway is a promising target for therapies. Currently, MET-specific therapies include ongoing clinical trials (NCT01744652, NCT01548144) that are combining immunotherapy and chemotherapy.

The DNA topoisomerase 2-alpha (TOP2A) is involved in the cell cycle, and it is crucial to cancer development as indicated by its expression, described as upregulated in several types of cancer, including pancreatic cancer [60, 61]. Network analyses identified TOP2A among the hub proteins in the protein–protein interaction network in glioblastoma, rectal adenocarcinoma, and endometrial cancer [60–62]. In pancreatic cancer, TOP2A is associated with tumor size, metastasis, tumor stage, poor prognosis. It is critical in pancreatic cancer development as shown by TOP2A knockout, suppressing proliferation, and migration/invasion in pancreatic cancer cells [63].

In addition to our validation, the CG expression values were tested in an independent dataset (GSE16515). The PCA and the heatmap hierarchical clustering analysis indicated that CG plays a central process in PDAC and is capable of classifying the data in two groups of normal and tumoral samples.

We used five genes with higher mRNA and protein expression according to immunohistochemical data to develop an ANN sample classifier based on gene expression. We achieve sensitivity and specificity of 0.97 and 0.88 respectively applying our ANN classifier in the test set. The development of automatic classifiers based on artificial intelligence could assist in the PDAC diagnosis. It was already pointed five possible PDAC biomarkers (FAIM3, IRANK3, DENND2D, PLBD1, AGPAT) based on gene expression, achieving a combined sensitivity of 100% and specificity of 94% respectively [64]; however, no automatic classification was produced. These five genes were pointed as potential biomarkers in PDAC diagnosis. Here, we not only pointed five genes independently differentially expressed among datasets but also created an automatic tool to classify the samples and give the probability of being normal or PDAC. In contrast with the list of five differentially genes reported by Irigoyen et al. 2018 [64], the CG list reported here did not include any of these genes.

The use of artificial intelligence to classify samples using PDAC genes expression was developed using support vector machines (SVM) and gene expression information of five genes (TMPRSS4, AHNAK2, POSTN, ECT2, and SERPINB5); although using different genes our ANN has better results compared with PDAC SVM classifier that showed on average for validation dataset sensitivity 94% and specificity of 86.67% [16]. The datasets used in both works are different, with this in mind, sample preparation or microarray technologies (Affymetrix and Illumina) could be possible explanations to different gene lists. Furthermore, the use of six datasets here in contrast with two datasets by Irigoyen et al. 2018 [64] could also produce different results. Another explanation for these gene list differences presented here could be due to PDAC subtypes already study in the gene expression and clinical level [10].

## Conclusions

The results indicate that PDAC presents a 60-core genes signature, with 58 genes upregulated and two downregulated. Among these upregulated genes, many are related to cell adhesion, migration, and extracellular matrix-receptor interaction; the two downregulated genes are associated with pancreatic functions. Immunohistochemistry analyses confirm the overexpression at the protein level of the 31 genes, validating our analysis. The five most overexpressed genes were related to tumor differentiation, cell migration, and metastasis. The PDAC-ANN trained using gene expression information could classify the samples in normal and PDAC with an f1-score of 0.94 and sensitivity = 0.97. The PDAC-ANN tool can only be used when the gene expression information from KRT19, LAMC2, MELK, MET, and TOP2A are available, in addition to min-max gene expression values rescaling. The PDAC-ANN is a free tool (Additional file 4) that can support in the pancreatic ductal adenocarcinoma diagnosis.

## Supporting information

Additional file 1: Table S1. Microarray gene expression results from the six studies here.

Additional file 2: Figure S1. PCA and hierarchical analysis of the CG expression values from GSE16515.

Additional file 3: Table S2. Excel file with links to protein expression level.

Additional file 4: Pancreatic ductal adenocarcinoma artificial neural network (PDAC-ANN) software.

## Availability and requirements

Project name: Pancreatic ductal adenocarcinoma artificial neural network (PDAC-ANN)

Project home page: https://github.com/freitasleandro/PDAC-ANN

Operating system(s): e.g. Platform independent

Programming language: Python 3.7

Other requirements: pandas, numpy, sklearn, keras, tensorflow, argparse.

License: e.g. GNU GPL v3.0.

Any restrictions to use by non-academics: license needed

## List of abbreviations

AI: artificial intelligence
ANN: artificial neural network
CG: core-genes
DEG: differentially expressed genes
ECM: extracellular matrix
GEO: Gene Expression Omnibus
mRNA: messenger RNA
PCA: principal component analysis
PDAC: pancreatic ductal adenocarcinoma
SVM: support vector machines
THPA: The Human Protein Atlas

## ADDITIONAL FILES

**Additional file 1: Table S1. Microarray gene expression results from the six studies here.** Excel file with information of spot ID, adjusted p-value ≤ 0.05, |logFC| ≥ 1,Gene symbol, Gene.ID (ENTREZ_GENE_ID).

**Additional file 2: Figure S1. PCA and hierarchical analysis of the CG expression values from GSE16515**. (a) The CG could produce similar results in both PCA and (b) heatmap hierarchical clustering analysis. The CG can classify the data into two groups of normal and tumoral samples.

**Additional file 3: Table S2. Excel file with links to protein expression level from the CG and links to immunohistochemical images available in THPA.**

**Additional file 4: Pancreatic ductal adenocarcinoma artificial neural network (PDAC-ANN) software.**

