## Supplementary figures and images for "PDAC-ANN: an artificial neural network to predict Pancreatic Ductal Adenocarcinoma based on gene expression"

### Additional file 2: Figure S1. PCA and hierarchical analysis of the CG expression values from GSE16515.

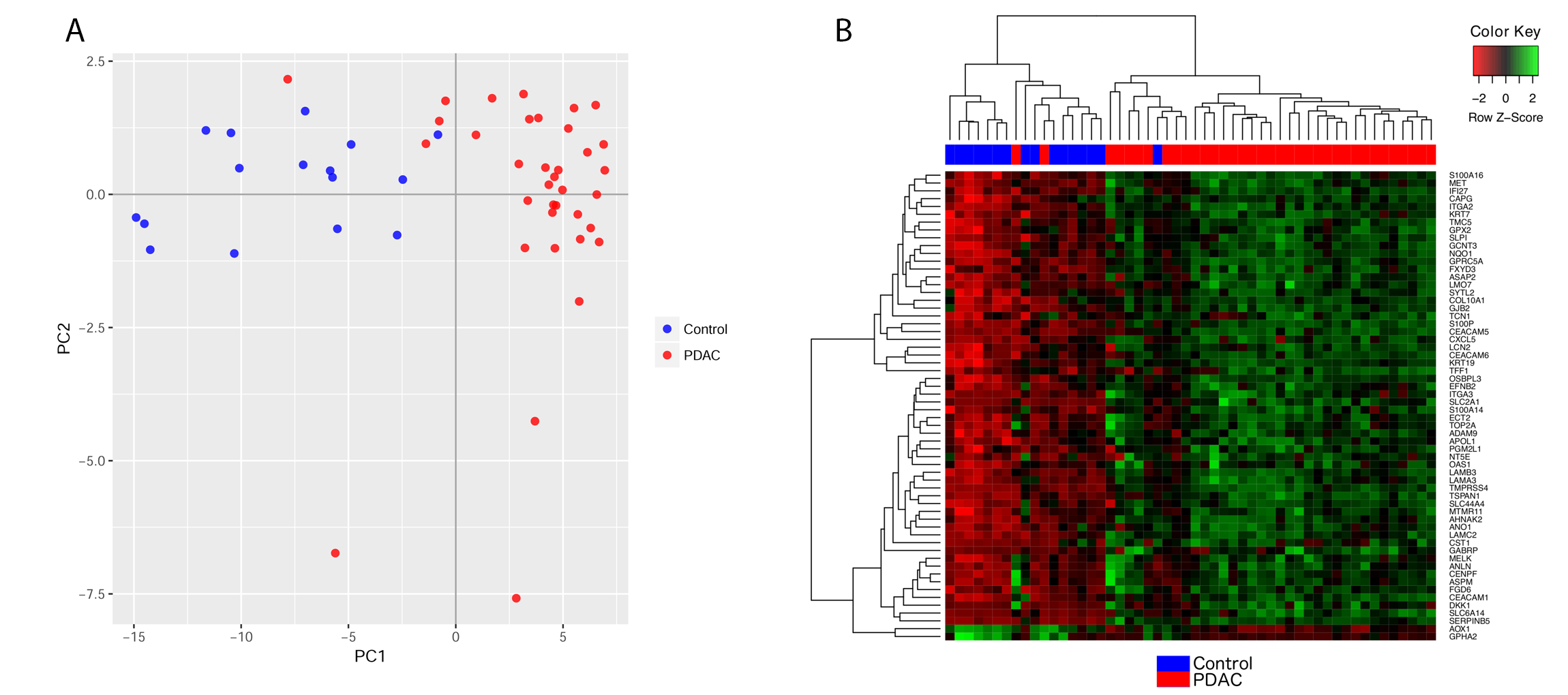
