## Additional file 4: Pancreatic ductal adenocarcinoma artificial neural network (PDAC-ANN) software. for "PDAC-ANN: an artificial neural network to predict Pancreatic Ductal Adenocarcinoma based on gene expression": PDAC.py

```
# coding: utf-8

#packages required
import argparse #deal with the parameters files https://www.cyberciti.biz/faq/python-command-line-arguments-argv-example/

#print("\n%s" % ("Welcome to pancreatic ductal adenocarcinoma sample classification.\nPDAC sample classification predict the sample status based on gene expression.\nThe genes used to classifiy the samples are:\nKeratin 19 (KRT19)\nLaminin Subunit Gamma 2(LAMC2)\nMaternal Embryonic Leucine Zipper Kinase (MELK)\nMET Proto-Oncogene, Receptor Tyrosine Kinase (MET)\nDNA Topoisomerase II Alpha(TOP2A)\n"))


__author__ = 'Dr. Leandro Martins de Freitas\nemail:' 
parser = argparse.ArgumentParser(description="Welcome to pancreatic ductal adenocarcinoma sample classification.\nPDAC sample classification predict the sample status based on gene expression.\nThe genes used to classifiy the samples are:\nKeratin 19 (KRT19)\nLaminin Subunit Gamma 2(LAMC2)\nMaternal Embryonic Leucine Zipper Kinase (MELK)\nMET Proto-Oncogene, Receptor Tyrosine Kinase (MET)\nDNA Topoisomerase II Alpha(TOP2A)\n")
parser.add_argument('-i','--input', help='Input file name',required=True)
parser.add_argument('-o','--output',help='Output file name', required=True)
#https://stackoverflow.com/questions/15301147/python-argparse-default-value-or-specified-value
parser.add_argument('-t','--threshold',help='PDAC threshold', nargs='?', const=0.78, type=float,default=0.78, required=False)
#parser.add_argument('-t','--threshold',help='PDAC threshold', required=True)
args = parser.parse_args() ## show values ##
print ("Input file: %s" % args.input )
print ("Output file: %s" % args.output )
print ("PDAC threshold: %s" % args.threshold )

file_name = args.input
output_file = args.output
threshold_value = args.threshold


import pandas as pd
import numpy as np
#import sys # deal with file name in the command line
from sklearn import preprocessing
from keras.models import load_model #load the ANN model


# load epitope training dataset
expression_data = pd.read_csv(file_name,  index_col='Samples',delimiter=";")

Samples = expression_data.index
#type(Samples)

# Returns a compiled model identical to the previous one
model = load_model('model/PDAC.h5')

prediction = model.predict(expression_data)
#prediction_class_binary = model.predict_classes(expression_data)


le = preprocessing.LabelEncoder()
le.fit(["NORMAL", "PDAC"])

prediction_binary = prediction[:,1]  > threshold_value

prediction_binary

len(prediction_binary)
prediction_binary_class = prediction_binary.astype(int)

prediction_binary_class

prediction_class_labels = list(le.inverse_transform(prediction_binary_class))

prediction_class_labels


#samples_validacao = pd.read_csv("/Users/leandromartinsdefreitas/Dropbox/PESQUISA/PALLOMA/topTable/samples_validacao_PDAC_0_1.csv", delimiter=",")
#samples_validacao = samples_validacao.transpose()
#samples_validacao.head()
#samples_validacao.values
#teste = le.inverse_transform(samples_validacao.values)

#teste.shape

#teste2 = teste.tolist()


# In[225]:


#y = np.hstack(teste2)
#print(y)


#Creating pandas dataframe from numpy array
prediction_df = pd.DataFrame({"Samples": Samples,'pr(Normal|data)':prediction[:,0],'pr(PDAC|data)':prediction[:,1],'Class':prediction_class_labels}) #, 'TRUE': y})
#print(prediction_df)


# In[240]:


prediction_df.to_csv(output_file, sep=';')
```
