## Additional file 4: Pancreatic ductal adenocarcinoma artificial neural network (PDAC-ANN) software. for "PDAC-ANN: an artificial neural network to predict Pancreatic Ductal Adenocarcinoma based on gene expression": README.txt

The script runs a deep learning model (PDAC-ANN) to classify samples in normal or pancreatic ductal adenocarcinoma (PDAC) using gene expression information. The genes use the samples prediction are Keratin 19 (KRT19), Laminin Subunit Gamma 2(LAMC2), Maternal Embryonic Leucine Zipper Kinase (MELK), MET Proto-Oncogene, Receptor Tyrosine Kinase (MET), DNA Topoisomerase II Alpha(TOP2A).

This initiative was built by 
Leandro Martins de Freitas -
Palloma Porto Almeida -
Cristina Padre Cardoso

The computer must have python 3.6 or higher to run the script and the packages:
pandas
numpy
sklearn
keras
tensorflow.

Help
python PDAC.py -h
help instructions to run the script.

Running the script
python script.py -i input_file.csv -o output_file.csv -t 0.78
-i input file
-o output file
-t threshold value to be considered PDAC sample  (0.78 by default)

Test run
python PDAC.py -i test/input_file.csv -o test/output_file.csv

The output file contains four columns. 
First column: samples name
Second column: the probability of the sample to be normal
Third column: the likelihood of the sample to be PDAC
Fourth column: the classification of the sample based on the threshold value.

The input file must be a csv file separated by ;
Fist line must be  the header
Example of an input csv file:
Samples;KRT19;LAMC2;MELK;MET;TOP2A
Sample1;0.84;0.78;0.58;0.55;0.50
Sample2;0.90;0.87;0.80;0.521;0.588
Sample3;0.923;0.908;0.86;0.53;0.68
Sample4;0.503;0.208;0.303;0.119;0.277
Sample5;0.59;0.181;0.35;0.41;0.56
